# Language models enable zero-shot prediction of RNA secondary structure including pseudoknots

**DOI:** 10.1101/2024.01.27.577533

**Authors:** Tiansu Gong, Dongbo Bu

## Abstract

Current deep learning-based models for predicting RNA secondary structures face challenges in achieving high generalization ability. At the same time, a vast repository of unlabeled non-coding RNA (ncRNA) sequences remains untapped for structure prediction tasks. To address this challenge, we trained RNA-km, a foundation language model that enables zero-shot prediction of RNA secondary structures including pseudoknots. For the end, we incorporated specific modifications into the language model training process, including k-mer masking strategy and relative positional encoding. RNA-km are trained on 23 million ncRNA sequences in a self-supervised manner, gaining the advantages of high generalization ability. For a target RNA sequence, we make a zero-shot secondary structure prediction with the attention maps provided by RNA-km and a specified minimum-cost flow algorithm. Our results on popular benchmark datasets demonstrate that RNA-km exhibits high generalization abilities, excelling in zero-shot predictions for RNA secondary structures. In addition, the attention maps provided by the model capture intricate structural relationships, as evidenced by accurate pseudoknot predictions and precise identification of longdistance base pairs. We anticipate that RNA-km enhances the predictive capacity and robustness of existing models, thereby improving their ability to accurately predict structures for novel RNA sequences.

## 1 Introduction

Ribonucleic acid (RNA) are polymer molecules with indispensable roles across a large variety of biological processes [1, 2], including transcription, translation [3], catalysis [4], gene expression regulation [5], protein synthesis [6], and degradation [7]. Approximately 5% of all RNA transcripts function as messenger RNAs (mRNAs), responsible for encoding proteins, while the substantial majority constitutes non-coding RNAs (ncRNAs)[8, 9]. Non-coding RNAs, a varied category that includes small nuclear ribonucleoproteins (snRNPs), small nucleolar ribonucleoproteins (snoRNPs), ribosomes, microRNAs, long ncRNAs, and telomerase, typically fold into specific structures and form stable RNA-protein complexes through interactions with proteins, carrying out crucial biological functions[10, 11]. These structures, coupled with RNA primary sequences, largely dictate the biological functions of RNAs [12], emphasizing the paramount importance of a comprehensive understanding of RNA structures.

RNA structures can be experimentally determined using X-ray crystallography [13], nuclear magnetic resonance [14], or cryo-electron microscopy [15]. Additionally, the investigation of RNA secondary structures, which often form through base pairing with hydrogen bonds and are stable and accessible within cells[16, 17], can be achieved experimentally through enzymatic and chemical probing methods [18, 19]. While these experimental technologies have made significant strides, their high associated costs [20] limit their widespread application. Despite the sequencing and collection of over 24 million ncRNAs in the RNAcentral database [21], only a fraction of them have experimentally determined structures [22]. Compared with these experimental determination technologies, computational prediction of RNA structures purely from RNA sequences is substantially efficient and has become a promising method for understanding RNA structures.

RNA secondary structure prediction has received extensive studies, and the existing methods can be divided into three categories: alignment-based methods, thermodynamic methods, and deep learning (DL)-based methods. Alignment-based approaches, built upon comparative sequence analysis, leverage sufficient homologous sequences and their alignments to achieve success in predictions [23, 24]. However, challenges arise from the limited number of known RNA families, as Rfam encompasses only several thousand such families [25]. The comparatively lower conservation of RNA, in contrast to proteins, also poses a theoretical constraint on the further improvement of alignment-based methods relative to the extensive repertoire of protein families.

Thermodynamic models quantify the stability of an RNA structure by calculating folding free energy change and then select the structure with the lowest free energy or the maximum expected accuracy as the most probable one in the entire structure ensemble [26, 27]. The widely used thermodynamic models incorporate a large volume of folding features (together with standard base pairs) to estimate the model parameters, and utilize dynamic programming to determine the final structure. Representative prediction approaches includes Vienna RNA [28], Mfold [29], LinearFold [30], and RNAstructure [31]. Despite subsequent enhancements through the consideration of local structural parameters, such as experimental factors (e.g., RNAstructure [31], RNAfold [28], RNAshapes [32]) and machine learning parameters (e.g., ContextFold [33], CONTRAfold [34], CentroidFold [35]), the overall performance of these thermodynamic-based methods seems to have reached a plateau. Notably, these methods often fall short in considering all base pairs obtained from tertiary interactions [36, 37], potentially overlooking crucial information in their predictions.

Recognizing the limitations of alignment-based and thermodynamic methods, de novo deep learning (DL) approaches have emerged as powerful tools to enhance RNA secondary structure predictions. Liberated from the constraints of physical laws or co-evolutionary constraints, these DL methods leverage the increasing repositories of RNA secondary structures. Models such as SPOTRNA [38], E2Efold [39], MXfold2 [40], and UFold [41] have demonstrated substantial improvements. Nevertheless, challenges persist in their low generalization capacity, hindering robust performance across diverse sequences and families [42, 43].

A major reason for the low generalization capacity of DL models is the limited availability of RNA secondary structure data. Unlike protein structure prediction, predicting RNA structures faces a systemic challenge due to the scarcity of training data: on one hand, the dataset size of known RNA secondary structures is limited — one of the most standardized and publicly available benchmarks, TR0 [38], contains only 10,814 RNA structures. On the other hand, the quality of known RNA secondary structures is also limited as some of them were obtained through comparative analysis, thus often leading to considerable deviation from their ground-truth structures [25, 44].

Recent advancements in self-supervised learning for RNAs highlight the potential of leveraging unlabeled data. In contrast to secondary structures, unlabeled RNA sequence data is abundant, with RNAcentral [21] housing over 23 million ncRNA sequences. Self-supervised models, such as RNA-FM [45], RNA-MSM [46], and ATOM-1 [47], extract meaningful RNA representations from these unannotated data, showcasing improvements in multiple downstream applications, including RNA secondary structure prediction. However, the extent to which unlabeled data can elevate the generality of RNA secondary structure prediction remains an open question.

In this work, we present RNA-km, an innovative RNA foundation model crafted through self-supervised training on a vast dataset comprising over 23 million RNA sequences from RNAcentral. The introduction of RNA-km offers the following key advancements:

- An effective RNA foundation model: We introduce RNA-km as an innovative RNA foundation model, providing meaningful RNA representations. RNA-km incorporates specific modifications, including k-mer masking and relative positional encoding, into the language model training process, thereby enhancing its efficacy.
- High generalization ability: RNA-km showcases robust generalization capabilities, particularly highlighted through its zero-shot prediction performance for RNA secondary structure.
- The ability to capture intricate structural motifs: The attention maps generated by RNA-km capture intricate structural relationships, evidenced by the model’s accurate pseudoknot predictions and precise identification of long-range interactions.

## 2 Methods

### 2.1 The RNA-km language model

RNA-km calculates RNA representations using a BERT-style encoder-only architecture. BERT [48] is a transformer-based contextualized language representation model to develop generalized understandings from extensive unlabeled data. Its training objective, predicting masked tokens from contexts, requires the model to learn complex internal representations of its input. These representations further enhance the performance of downstream tasks. Early studies have highlighted the significance of these representations for RNA, showcasing capabilities in tasks such as secondary structure prediction and binding site prediction [45].

As depicted in Figure 1, RNA-km takes an RNA sequence *x* with *L* bases as its input. Initially, each base type (A, C, G, U, or N) is converted into a learnable embedding of *D* dimensions using an embedding layer. Subsequently, RNA-km captures contextual information from the input embedding (the sum of the base embedding and a relative positional encoding) through a specially designed model with an encoder-only transformer architecture [49]. The transformer-based model comprises 12 transformer-based bidirectional encoder layers. Each transformer encoder layer includes a multi-head self-attention module and a position-wise feed-forward network. The output embedding is the sequence representation forming an array of size *L × D*.

**Fig. 1.**
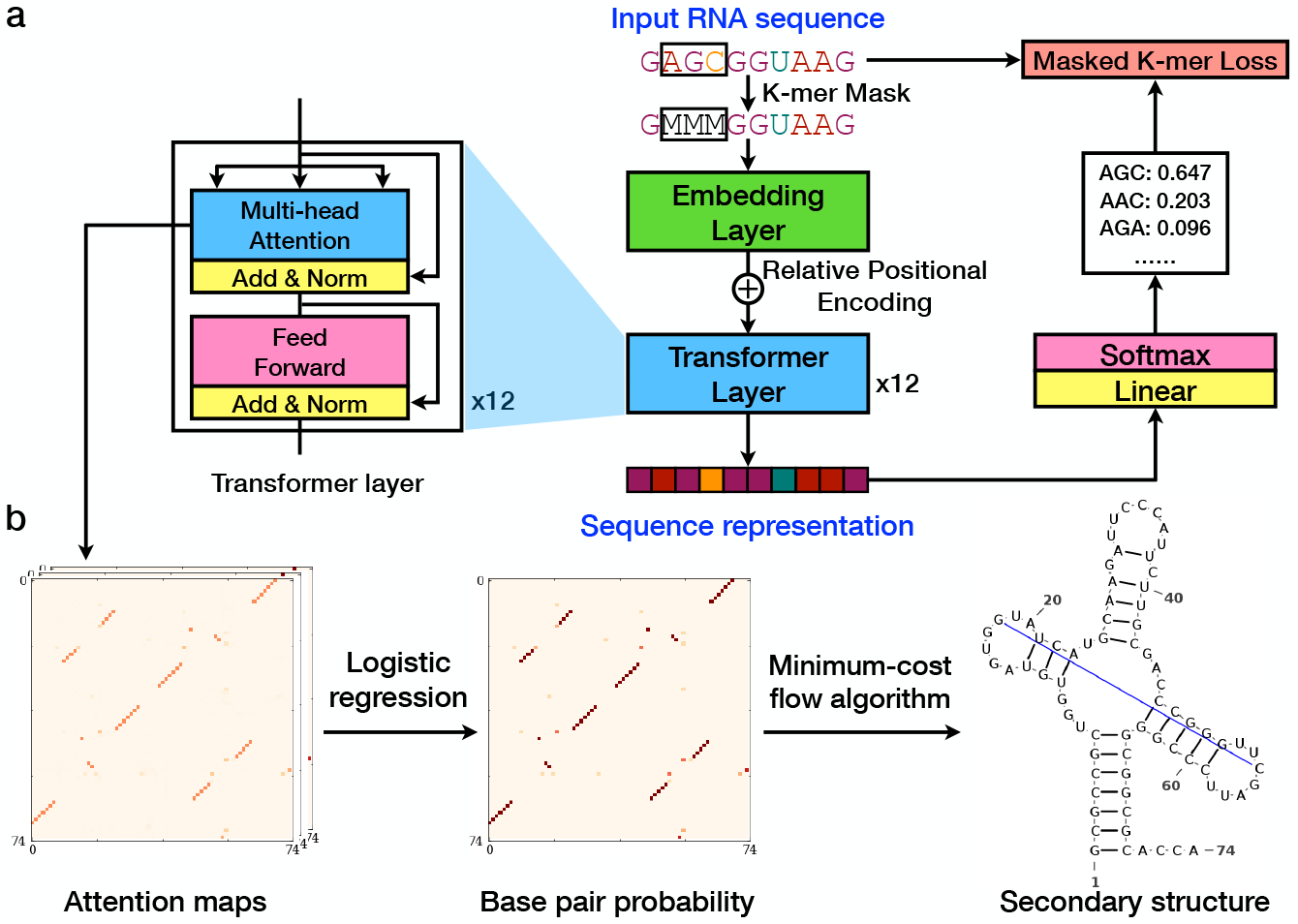
Overview of the self-supervised training process and zero-shot secondary structure prediction of RNA-km. **a**. The transformer architecture of RNA-km and its pretraining process. A k-mer masking strategy and relative positional encoding are employed, exemplified here with *k* = 3: the three bases within the black frame ‘‘`AGC`” were masked, denoted as ‘‘`MMM`”. **b**. Predicting RNA secondary structure from the attention maps acquired by RNA-km. For a target RNA sequence, RNA-km provides the attention maps, based on which we estimate base pair probabilities. Subsequently, these probabilities are utilized to predict RNA secondary structure through a minimum-cost flow algorithm. The attention maps are unexposed to known RNA structures, rendering the entire process a zero-shot prediction and thus showing the advantages of high generalization ability. In addition, the use of minimum-cost flow algorithm gains the ability to predict secondary structures even with pseudoknots (shown as blue line)

Positional encoding is incorporated into the input embedding at the base of the encoder stack. The original transformer architecture employed absolute sinusoidal positional encoding to convey information about token positions to the model. In RNA-km, we substitute the absolute sinusoidal positional encoding with a relative one [50]. Unlike absolute encoding methods, which assign a unique fixed vector based on sinusoidal functions and struggle to extrapolate beyond the trained context window, our relative positional encoding enables the model to enhance local context awareness and exhibits reduced sensitivity to variations in sequence length. In addition, the missing of the initial or final segments of an RNA sequence will have little impact on the intrinsic relationships.

RNA-km is pre-trained with a typical Masked Language Modeling (MLM) loss:

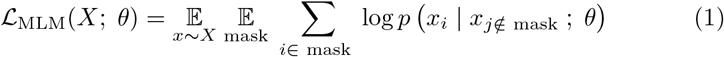

### 2.2 Pre-training process

We represent an RNA sequence as a string of nucleotide bases, including A, C, G, U, or N (representing the bases other than the four bases). Each nucleotide base is considered as an individual token. In the pre-training stage, we randomly mask 15% of the bases within a given RNA sequence and task our model, RNA-km, with predicting the missing bases in a self-supervised manner. To enhance RNA-km to capture local dependencies, we employ a mask strategy that 80% of the masks are contiguous spans of length *k* (i.e., k-mers, where 3*≤ k≤* 8) rather than a single base.

RNA-km was trained on a dataset comprising over 23 million ncRNA sequences sourced from RNAcentral version 22.0 (https://ftp.ebi.ac.uk/pub/databases/RNAcentral/) [21]. To accommodate memory constraints during training, we crop RNA sequences to nonoverlapping fragments of 512 bases when their original length exceeds this threshold [51]. Subsequently, we employ MMseqs2 [52] to cluster the fragments at a sequence identity cut-off of 80%, resulting in 8.9 million clusters. From these clusters, we randomly select 10,000 clusters to form a held-out validation set. During the training process, each cluster is sampled with equal probability.

For the balance of performance and model size, we set the embedding dimension *D* as 1024, and used 12 transformer encoder layers with 16 attention heads in the study, resulting in a total of 152 million parameters. We train RNA-km with a batch size of 1 million tokens for 600,000 steps on GPU (Tesla V100 PCIe, 16GB memory).

### 2.3 Zero-shot secondary structure prediction

#### 2.3.1 Deriving the base pairing probability from attention maps

Zero-shot transfer learning is defined as deploying a model on a new task without additional supervision for task-specific adaptation [53, 54]. Language models, served as general-purpose models, can be trained once and then applied to a variety of possible tasks. Zero-shot prediction provides a systematic way to assess the generalization capabilities of pretrained models [55].

The zero-shot secondary structure prediction setting in this work follows Rao *et al* [56], which showed that transformer language models trained on large databases of protein sequences learn to predict protein contacts in their attention maps with little supervision. We exploit this observation to measure RNA language model’s knowledge of secondary structure. Specifically, we employ a logistic regression to the attention maps of RNA-km. Let *bp*_*ij*_ be a boolean random variable which is true if bases *i, j* form a base pair and false otherwise. For a transformer-based language model with *M* layers and *H* attention heads per layer, let *A*_*mh*_ be the *L × L* symmetrized and APC-corrected [57] attention map for the *h*-th head in the *m*-th layer, and 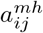 be the value of the attention map at position *i, j*. Then the probability of a base pair *P* (*bp*_*ij*_) is calculated as:

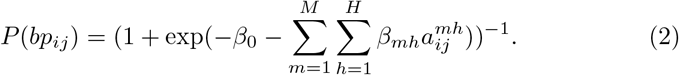

The parameters *β* are fit in scikit-learn [58] using L1-regularized logistic regression with regularization weight *λ* = 0.15. The regression is fit using 5 RNA randomly selected from a subset of PDB containing 532 RNAs.

To assess the potential bias from the chosen 5 RNAs, we conducted a variability analysis through 20 bootstrapped samples, each comprising 5 training RNAs randomly selected from the complete set of 532 RNAs. The average F1 score for the secondary structure prediction by RNA-km was 0.628, with a standard deviation of 0.00864. We also performed experiments using data comprising more than 5 RNAs, but observed no significant performance change. These findings imply that the selection and fitting of a subset of RNAs primarily serves as a dimension reduction method, exerting minimal influence on the zero-shot prediction performance of RNA-km.

#### 2.3.2 Secondary structure construction based on generated base pair probability

We construct a secondary structure from this *L × L* base pairing probability matrix of *P* (*bp*_*ij*_) using a minimum-cost flow algorithm proposed in KnotFold [59]. Specifically, we first calculate an RNA-specific structural potential by treating *P* (*bp*_*ij*_) as the prior probability and using a constant reference probability 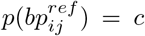. For an RNA with secondary structure *S* (can be regarded as the set of (i,j) where *bp*_*ij*_ = 1), the structural potential is formally described as:

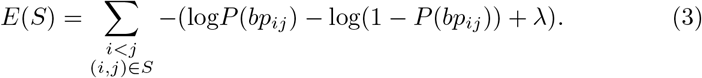

The parameter *λ* is determined using 5 RNAs randomly sampled from the 532 RNAs extracted from PDB, the same process as zero-shot experiment setting. Then we apply a minimum-cost flow algorithm proposed in KnotFold [59] to obtain the secondary structure with the lowest structural potential.

### 2.4 Datasets

In this study, we assessed the performance of various prediction approaches using RNA extracted from several databases. Our evaluation incorporated the following datasets:

i. Protein Data Bank (PDB): PDB [22] serves as a repository for highresolution RNA/DNA tertiary structures, elucidated through experimental techniques such as X-ray crystallography, nuclear magnetic resonance, and cryo-electron microscopy. We collected the RNAs with 3D structures from PDB, and removed the RNA structures if they were primarily determined by protein or DNA binding partners. We further removed the redundant sequences by running CD-HIT-EST [60] with a cut-off threshold of 80% sequence identity. This way, we obtained a dataset comprising 532 RNAs. Subsequently, the secondary structure of these RNAs was determined using DSSR [61]. Out of the 532 RNAs, 158 are pseudoknotted. This dataset was employed for benchmarking the performance of zero-shot RNA secondary structure prediction, as well as for fine-tuning pretrained models.
ii. bpRNA-1m: The bpRNA-1m dataset [44] is a comprehensive dataset comprising 102,318 RNA secondary structures collected from various sources. We utilize, TR0, a subset of bpRNA-1m extracted by SPOTRNA [38] to benchmark the zero-shot RNA secondary structure prediction performance of self-supervised models. TR0 contains 10,814 RNAs, among which 999 have pseudoknots.

## 3 Results

### 3.1 Decoding semantical information in RNA sequences with RNA-km

To investigate the semantical information learned by RNAnt, we examine the sequence representations of RNAs from RNAcentral [21]. Utilizing t-SNE [62], we project these representations onto a 2-D plane. Figure 2a illustrates that RNA-km captures semantics related to RNA types. Sequences with the same RNA types exhibit proximal representations, while those with different types manifest more distinct representations. This outcome shows RNA-km’s proficiency in acquiring semantically rich representations of RNA sequences.

**Fig. 2.**
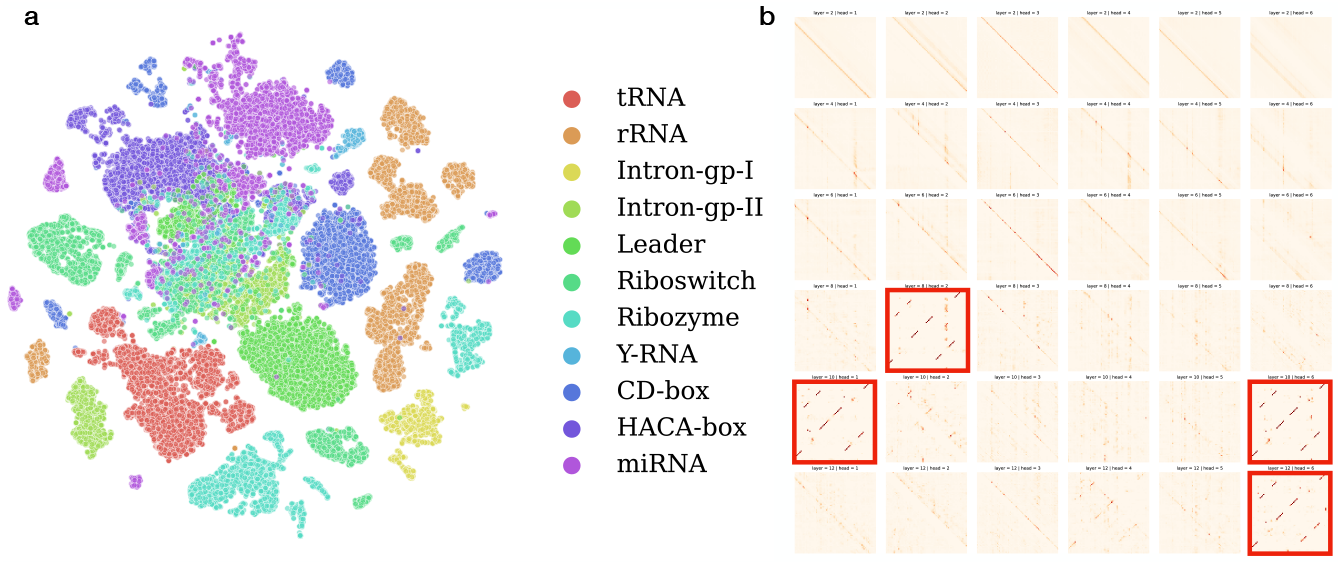
RNA-km learns semantically rich representation of sequences. **a**. t-SNE visualization of the sequence representations generated by RNA-km, color-coded according to RNA types. **b**. Depiction of attention maps acquired from the first six attention heads of the even layers (PDB entry: `5E6M`). Certain attention maps enable direct prediction of RNA base pairing (highlighted by red frames), achieving an F1 score exceeding 0.4 (See Supplementary Fig. 1 for a high-resolution version).

To further explore the model’s capability in capturing base pairing relationships, we analyze the attention maps obtained from RNA-km, as depicted in Figure 2b. Specifically, the attention maps of 5E6M from the initial six attention heads within the even layers are demonstrated. We observe that the first half of the transformer layers excels in extracting local features and low-dimensional representations. Conversely, the subsequent six layers demonstrate a preference in capturing high-level representations, including base pairing patterns.

Intriguingly, specific attention maps exhibit the capacity for direct prediction of base pairing, a binary classification task (forming base pair or not) on the upper triangle probability matrix. Specifically, we transfer the attention map into a binary matrix using a pre-defined threshold, and evaluate the performance of the attention map by F1 score on a set of 532 non-redundant RNAs collected from PDB [22] (see Methods for details of dataset). As highlighted in red frames in Figure 2b, specific attention maps yield F1 score of over 0.4. Notice the attention maps predicts base pairing in a zero-shot manner [53, 55]. This implies that RNA-km can discern base pairing relationships without any prior training examples on base pairing or secondary structures. Thus, our findings demonstrate RNA-km’s capability in capturing base pairing relationships, and it enables zero-shot prediction of RNA secondary structure.

### 3.2 RNA-km enables zero-shot prediction of RNA secondary structure prediction

To further evaluate the generalization ability of RNA-km, we conduct another zero-shot experiment by directly utilizing all the attention weights of the pretrained model to predict RNA secondary structures, characterized by hydrogen bonding patterns in canonical Watson-Crick or wobble base pairs [17].

Specifically, we extract base pairing probabilities from RNA-km’s attention maps using a logistic regression approach [56]. The secondary structure is subsequently constructed from these probabilities using a minimum-cost flow algorithm, a technique introduced in previous work, KnotFold [59]. Logistic regression parameters are fit using only 5 RNAs randomly selected from a set of 532 non-redundant, with the remaining 527 RNAs utilized for benchmarking the models’ performance (see Methods for details). To ensure a fair comparison in zero-shot secondary structure prediction, we apply the same process to the attention weights learned by RNA-FM [45]. The accuracy of the prediction is evaluated using precision, recall, and F1 score of the base pairs, averaged across the dataset.

As illustrated in Figure 3, RNA-km demonstrates a high F1 score of 0.627 on the PDB benchmark, showcasing its understanding of semantic relationships between base tokens. Notably, RNA-km, pretrained solely on RNA sequences to recover missing bases based on context tokens without encountering actual RNA secondary structures, exhibits commendable generalization capability. In comparison, RNA-FM yields an F1 score of 0.433. Head-to-head analysis suggests that in the majority of cases, RNA-km outperforms RNAFM. We observed that the cases where RNA-FM outperforms RNA-km mainly appear for the sequences shorter than 100 bases (94 out of 95), some of them are even shorter than 40 bases. The underlying reason might be that RNA-km were mainly pretrained on long RNA sequences.

**Fig. 3.**
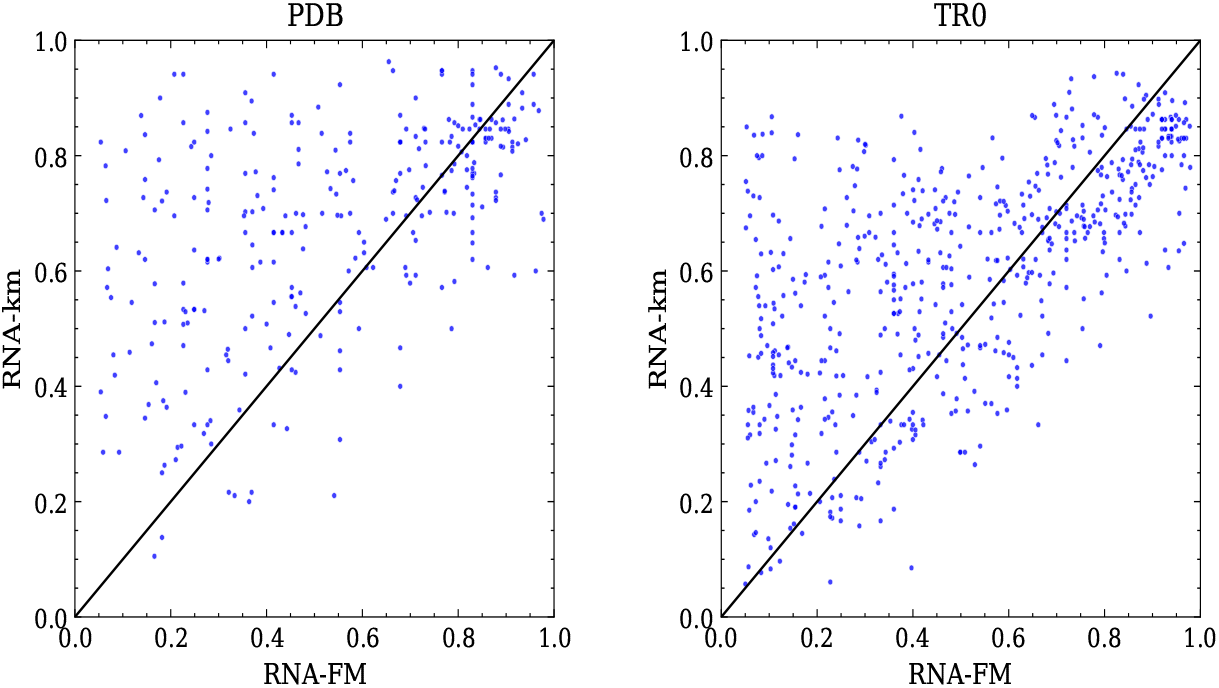
Performance of zero-shot prediction of RNA secondary structures by RNA-FM and RNA-km. Metric: F1 score; Datasets: PDB and TR0. Left panel: head-to-head comparison of the built structures of RNA-FM and RNA-km over a benchmark of 527 non-redundant proteins collected from PDB. RNA-FM and exhibited an overall F1 score of 0.433 and 0.627, respectively. Right panel: head-to-head comparison of the built structures of RNA-FM and RNA-km over TR0 [38], a dataset comprising 10,814 RNAs from bpRNA. RNA-FM and exhibited an overall F1 score of 0.339 and 0.456, respectively

Furthermore, we evaluate the performance of RNA-km on the SPOTRNA [38] training set TR0, a subset of bpRNA-1m [44] with 10,814 RNAs. As depicted in Figure 3, RNA-km demonstrates an F1 score of 0.456 on TR0, whereas RNA-FM yields an F1 score of 0.339. Both models exhibit a performance decrease compared to the PDB benchmark. Possible reasons lie in the greater diversity of structure patterns in TR0, and lower quality of secondary structure labels in TR0 [25, 44].

### 3.3 RNA-km identifies pseudoknots from sequences

Pseudoknots, intricate bipartite helical structures formed by pairing a singlestranded region inside a stem-loop structure with a complementary stretch outside [63], serve as crucial structural motifs for RNA stabilization and function [64, 65]. Despite their significance, accurate prediction of RNA secondary structures involving pseudoknots remains a challenging task due to its non-nested nature [66].

To assess the ability of pretrained models to identify RNA structural motifs including pseudoknots, we conducted experiments specifically focusing on pseudoknotted RNAs within the PDB (158 out of 527) and TR0 (999 out of 10,814) benchmarks. We also tested the five RNAs used to determine logistic regression parameters.

As depicted in Figure 4, RNA-km demonstrates an F1 score of 0.577 for all pseudoknotted RNAs on PDB and 0.527 on TR0. These results suggest that pretrained models successfully capture the relationship within the pseudoknots. Notably, the advantage of RNA-km over RNA-FM is more pronounced for pseudoknotted structures, with a performance difference of 0.199 on PDB and 0.274 on TR0, both exceeding the difference observed for all RNAs (0.194 on PDB, and 0.117 on TR0). Note that RNA-km and RNA-FM construct secondary structures using the same process with the only difference in the attention weights provided by the pretrained models. Thus, this performance clearly demonstrated that RNA-km’s pretrained model acquires more information about the structural relationships inherent in pseudoknots compared to RNA-FM.

**Fig. 4.**
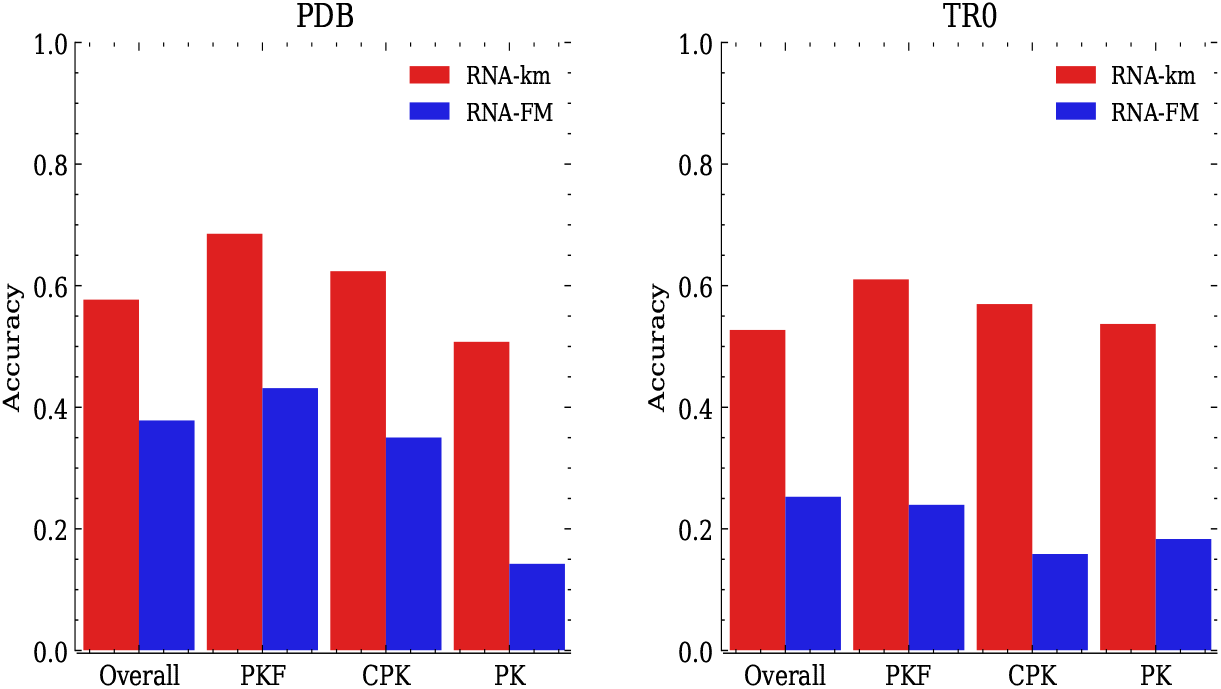
Performance of RNA-FM and RNA-km to secondary structure prediction for pseudoknotted RNAs in PDB and TR0 datasets. Left panel: Performance evaluated on pseudoknotted RNAs in PDB. Right panel: Performance evaluated on pseudoknotted RNAs in TR0. “Overall”, “PKF”, “CPK”, “PK” means the overall F1 scores of predicted structures, the prediction accuracy for pseudoknot-free base pairs, the prediction accuracy for crossing-pseudoknot base pairs, the prediction accuracy for pseudoknotted base pairs, respectively

To better evaluate the performance of both models on identifying pseudoknots, we perform a base-pair-level analysis similar to KnotFold [59]. Specifically, we classify base pairs in the target structures into three categories based on their structural motifs: (*i*) pseudoknot-free (PKF) base pairs, (*ii*) pseudoknotted (PK) base pairs, and (*iii*) crossing-pseudoknot (CPK) base pairs. CPK and PK base pairs, being non-nested and associating more closely with pseudoknot structures, carry more significance for pseudoknot identification. As summarized in Fig. 4, RNA-km gains a more pronounced advantage in predicting CPK and PK base pairs, highlighting its capability in capturing intricate pseudoknot structures.

In Figure 5, we present a concrete example: 5E6M is a 74-nucleotide-long RNA featuring four bulges together with a pseudoknot, connecting the 18th and 54th bases. Both RNA-FM and RNA-km accurately predict the secondary structure, achieving F1 scores of 0.842 and 0.930, respectively. However, a noteworthy difference emerges: RNA-FM mistakenly reports secondary structures with three bulges and fails to identify the pseudoknots while RNA-km successfully identifies both the four bulges and the pseudoknot. To explore the reasons behind these disparities, we examine the base pair probabilities. Comparative analysis with the ground truth reveals that the base pair probabilities reported by RNA-FM exhibit more noise and show a weak signal to the missing bulge and the pseudoknot. In contrast, RNA-km clearly reports the base pairing of them. This observation appears across multiple cases, demonstrating that the attention maps obtained by the pretrained model of RNA-km can more clearly delineate base-pairing relationships, especially for intricate structural motifs such as pseudoknots.

**Fig. 5.**
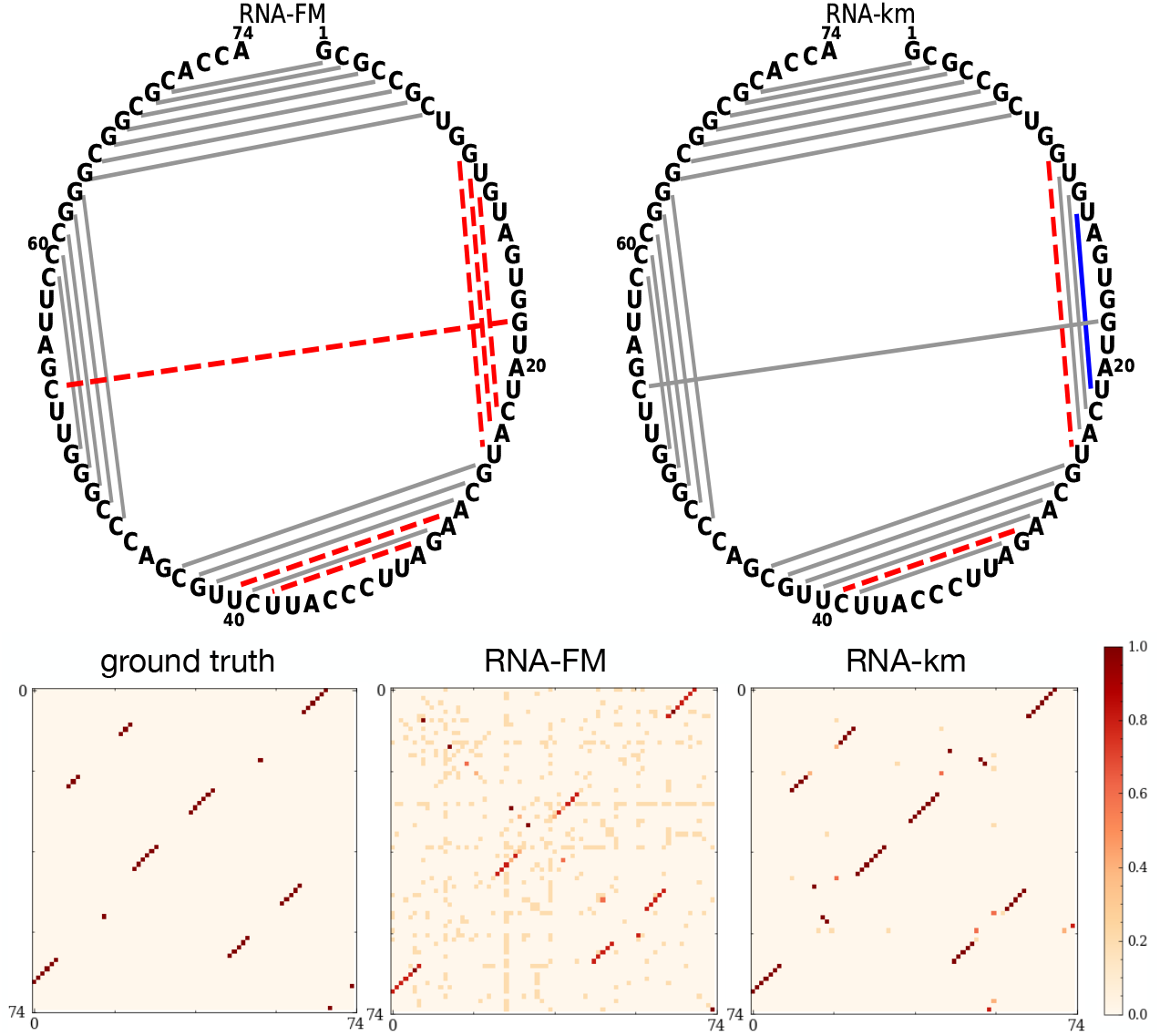
Predicting secondary structure of 5E6M using RNA-FM and RNA-km. Left panel: The predicted secondary structure by RNA-FM. The red dashed lines represent the missing base pairs, while the blue lines represent the false-positive base pairs. Right panel: The predicted secondary structure by RNA-km, in which pseudoknots were successfully predicted

### 3.4 RNA-km captures long-range interactions from sequences

Long-range interactions play an important role in understanding the folding patterns of RNA structures, as they often contribute to the stability and functionality of the molecule [67]. Accurately modeling these long-range relationships is crucial for precise RNA secondary structure prediction. Our intuition suggests that attention models, renowned for their adeptness in capturing contextual dependencies, are well-suited for modeling such long-range interactions. Thus, we assess the models’ performance in capturing long-range interactions.

Here, following the criteria outlined by Doshi et al. [67], we define “longdistance base pairs” as pairs separated by at least 101 bases in the RNA sequence. To evaluate the models’ performance in capturing these long-range interactions, we adopt the average accuracy for long-distance base pairs as our evaluation criteria.

As depicted in Figure 6, on the PDB benchmark, RNA-km achieves an accuracy of long-range interactions of 0.460, higher than RNA-FM’s accuracy of 0.207. A similar trend is observed on TR0, where RNA-km attains an accuracy of long-range interactions of 0.228, surpassing RNA-FM’s accuracy of 0.145. These results highlight RNA-km’s capacity in predicting long-distance base pairs, underscoring its effectiveness in capturing long-range interactions in RNA structures.

**Fig. 6.**
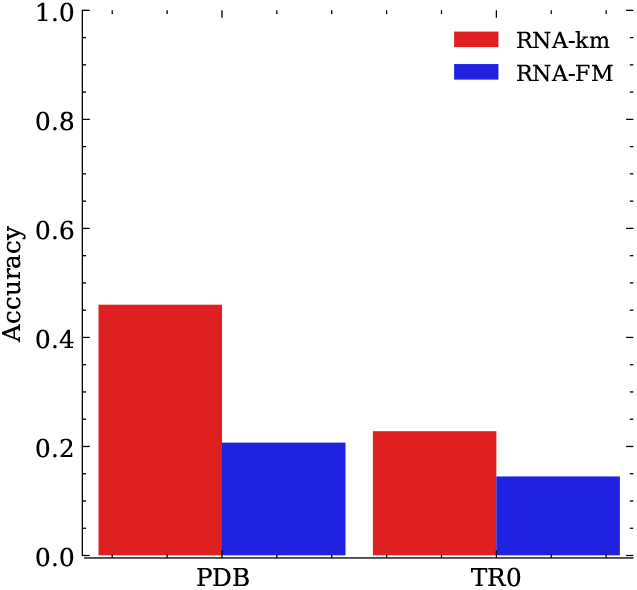
Performance of RNA-FM and RNA-km for long-range interactions in PDB and TR0 dataset.

## 4 Conclusion

In summary, we highlight RNA-km’s robust generalization capabilities on diverse benchmarks, particularly its advance in zero-shot predictions for RNA secondary structures. Through transfer learning on limited data, RNA-km achieves high prediction performance, surpassing current deep-learning models (See Supplementary Fig. 2). The attention maps derived from RNA-km adeptly capture intricate structural relationships, as evidenced by accurate pseudoknot predictions and precise identification of long-range interactions. Besides, we also demonstrate the power of k-mer masking and relative positional encoding through ablation study (see Supplementary Table 1, 2).

We observe a performance decrease of RNA-km for zero-shot secondary structure prediction on TR0 compared to the PDB benchmark. Possible reasons lie in the greater diversity of structure patterns in TR0, and lower quality of secondary structure labels in TR0. Our future work will involve enhancing the performance on these targets.

The integration of RNA-km into existing RNA secondary structure prediction frameworks holds significant promise. Incorporating RNA-km’s representations and attention weights as additional features have potential to enrich the predictive capacity and resilience of established models, enhancing their ability to predict structures for novel RNA sequences.

As a general-purpose language model, RNA-km could further extend its utility to various downstream tasks beyond secondary structure prediction. Exploring applications in tertiary structure prediction and investigating RNA function and disease associations present exciting avenues for future research. We anticipate that the understanding of RNA provided by RNA-km will greatly contribute to advancements in computational biology and RNA-focused research.

